# Boat anchors not OK: Loss of Dugong grass (*Halophila ovalis*) population structure in Havelock island of Andaman and Nicobar Islands, India

**DOI:** 10.1101/642579

**Authors:** A.K. Mishra, N.S. Sumantha, A. Deepak

## Abstract

Anthropogenic disturbance due to deployment of boat anchors and loss of seagrass ecosystem is not well understood in India. So, we used Govind Nagar beach of Havelock Island of Andaman and Nicobar Islands (ANI)to assess the impacts of boat anchors from traditional fishing and recreational activities on the seagrass *Halophila ovalis* population structure. *H. ovalis* density, biomass, morphometrics, canopy height and percentage cover were estimated from two stations of Govind Nagar beach i.e., one highly impacted from boat anchors (Station1) and a sheltered station (Station 2). A clear evidence in reduction of shoot density of *H. ovalis* was observed at station 1, exception was similar apex densities between both stations. *H. ovalis* morphometrics, such as number of leaves per shoot, leaf length, width and horizontal rhizome length were observed with significant lower values at station 1 compared to the sheltered station 2. Reduction in seagrass morphometrics also resulted in the loss of seagrass canopy height and percentage cover. A clear evidence of loss of seagrass population structure under the influence of physical disturbances caused by boat anchors were observed. We report for the first time the impacts of boat anchors on seagrass ecosystems of India and our results pitch for wider studies across India. The impact of boat anchors is small-scale, but in long-term loss of seagrass ecosystem services will have dire consequences on fish habitat and carbon storage. Therefore, proper management and conservation measures should be taken to prevent the loss of important dugong grass habitats of ANI.

**Highlights:**

- Physical disturbances caused by boat anchors decreased the shoot density of *H. ovalis* by 1.2-fold.
- 1 to 2-fold reduction in canopy height and the morphological features of individual plants were observed due to damage caused by boat anchors
- Habitat disturbance reduced 1.6-fold percentage cover of *H. ovalis* at Havelock Island of Andaman and Nicobar Islands, India

## Introduction

Seagrass ecosystems represent one of the richest and widely distributed coastal habitats in the ocean, that support a range of keystone and ecologically important marine species from all trophic levels (Short et al., 2011). These ecosystems provide 24 different types of ecosystem services greater than many terrestrial and marine habitats (Short et al., 2011; Nordlund et al., 2016) and contribute significantly to the health of coral reefs, mangroves and salt marshes (Unsworth et al., 2010). Seagrass ecosystems form important habitat and nurseries to1/5^th^ of 25 commercially important fish population and endangered sea cows and seahorses (Cullen-Unsworth et al., 2018) that directly support artisanal fisheries and the livelihoods of millions of coastal communities (Nordlund et al.,2017). They sequester 35 times faster and store more carbon (Duarte et al., 2013a) that helps in mitigation of climate change. Along with supporting fisheries and acting as carbon sink, seagrass meadows protect the shoreline (Boudouresque et al., 2016), diminishing wave energy and trapping sediments (Ondiviela et al., 2014), regulating nutrient cycling (Costanza et al., 2014) and acting as bioindicators of coastal pollution (Lewis and Devereaux, 2009). Though seagrass ecosystems provide valuable ecosystem services, they have received less attention than coral reefs and mangroves in terms of research, management and conservation practices (Nordlund et al., 2016). Seagrass ecosystems are declining globally around 7% yr^-1^ under the influence of anthropogenic pressure (Waycott et al., 2009; Lewis and Devereaux, 2009), which have led to extinction risk of 11 species of seagrass worldwide and three under endangered category (Short et al., 2011).The loss of biodiversity under the influence of anthropogenic pressure is pushing the ecosystem boundaries and biodiversity towards mass extinction worldwide (Intergovernmental Science-Policy Platform on Biodiversity and Ecosystem Services, (IPBES), 2019) which emphasizes the importance of biodiversity conservation from anthropogenic disturbances.

One of the major contributors of seagrass decline worldwide is coastal development and modification caused due to human settlement (Bjork et al., 2008), which has led to significant reduction of coastal water quality, nutrient enrichment leading to eutrophication (Unsworth et al.,2015; Maxwell et al., 2016), increased sedimentation from land run-off, tourism activities and destructive fishing practices (Dies, 2000; Duarte et al., 2004; Short et al., 2011). Both tourism and fishing activities indulge the use of various boats, which are parked in the shallow waters by deployment of boat anchors. Boat anchors are of serious concern (Okudan et al., 2013) which represents a long-term small-scale physical disturbance to shallow water seagrass ecosystems (Macreadie et al., 2015) leading to permanent damage of seagrass root and rhizome structure (Bourque et al., 2015) leading to loss of seagrass meadows. These loss of seagrass meadows due to boat anchors are reported worldwide for species like *Zostera marina* of San Francisco Bay, USA (Kelly et al., 2019) and Studland Bay, UK (Collins et al., 2010; Unsworth et al., 2017), *Posidonia oceanica* in Mediterranean Sea (Francour et al., 1999; Milazzo et al., 2004; Ceccherelli et al., 2007; Montefalcone et al., 2006, 2008), mixed seagrass species of Western Australia (Walker et al., 1989)and Rottnest Island, Australia (Serrano et al., 2016) and *Halodule wrightii* of Brazilian coast (Creed and Filho, 1999). Loss of these seagrass meadows resulted in eventually loss of valuable ecosystem services, such as release of stored carbon of 4.2 kg C_org_ m^-2^ (Serrano et al., 2016) and loss of fish habitats and herbivory for sea cows (Serrano et al., 2016; Unsworth et al., 2017).

India has an estimated cover of 517Km^2^ of seagrass beds consisting of 7 genera and 16 species (Patro et al., 2017; Thangaradjou and Bhatt, 2018) distributed along its coastline along with Andaman and Nicobar Islands (ANI) and Lakshadweep islands (Geevarghese et al., 2016; Ramesh et al., 2018). The seagrass *Halophila ovalis* has a pan India distribution and recorded around the east coast at Chilika lagoon, Odisha (Priyadarshini et al., 2014; Ganguly et al., 2018), Gulf of Mannar, Tamilnadu (Patro et al., 2017) and Andaman and Nicobar Islands (Ragavan et al., 2016). ANI has 13 seagrass species covering an area of 29.42 square Km (Nobi et al.,2013; Fortes et al., 2018) and distributed around mudflats and sandy regions from intertidal zone to 10-15m depth (Jagtap et al., 2003; Ragavan et al., 2016; Thangaradjou and Bhatt, 2018). *H. ovalis* has a frequent distribution around ANI mostly in the intertidal regions, where the plant is found in individual patches or mixed with other seagrass species such as *Halodule uninervis* and *Thalassia hemprichii* (Ragavan et al., 2016). *H. ovalis* is fastest growing seagrass species in these regions (Vermaat et al., 1995; Bharathi et al., 2014) and act as a preferred food source for the endangered *Dugong dugon* (Nakaoka and Aioi, 1999).

Tourism is a major source of income in Havelock islands (Swaraj Deep) of ANI because of its natural beaches and under water marine life, such as coral reefs and associated biodiversity. Being a tourist hotspot, these islands have seen a rapid increase in number of boats operating at this island for SCUBA diving, fishing (traditional and recreational) and various other recreational activities. Saying that, the impacts of these increased boat anchoring on seagrass species of ANI is not well understood. So, this proposed work will evaluate the density, biomass, morphometrics and canopy structure of *H. ovalis* meadows of Havelock islands under the influence of human activities such as boat anchoring to understand the population structure.

## Methods

Havelock island (11°58’ 12.27” N; 92°59’ 52.27” E) is situated in south east region of Andaman and Nicobar Islands (Fig.1). The tidal variation of Havelock island is between 0.01 to 2.45m, temperature ranges between 26.28 to31.67°C and the salinity range is between 32 to 35 ppt. Number of fishing and recreational boats anchored are about ∼120 and ∼20 respectively at Govind Nagar beach of Havelock island. Two stations with the Govind nagar beach was selected (Fig.1). The stations were within 500m far from each other and were separated by dead coral patches. The station one has more boat anchors deployed than the station two as it was more sheltered by dead coral patches (Fig.2c and d) and there is a chance of boat wreckage during high wave actions.

**Fig. 1.**
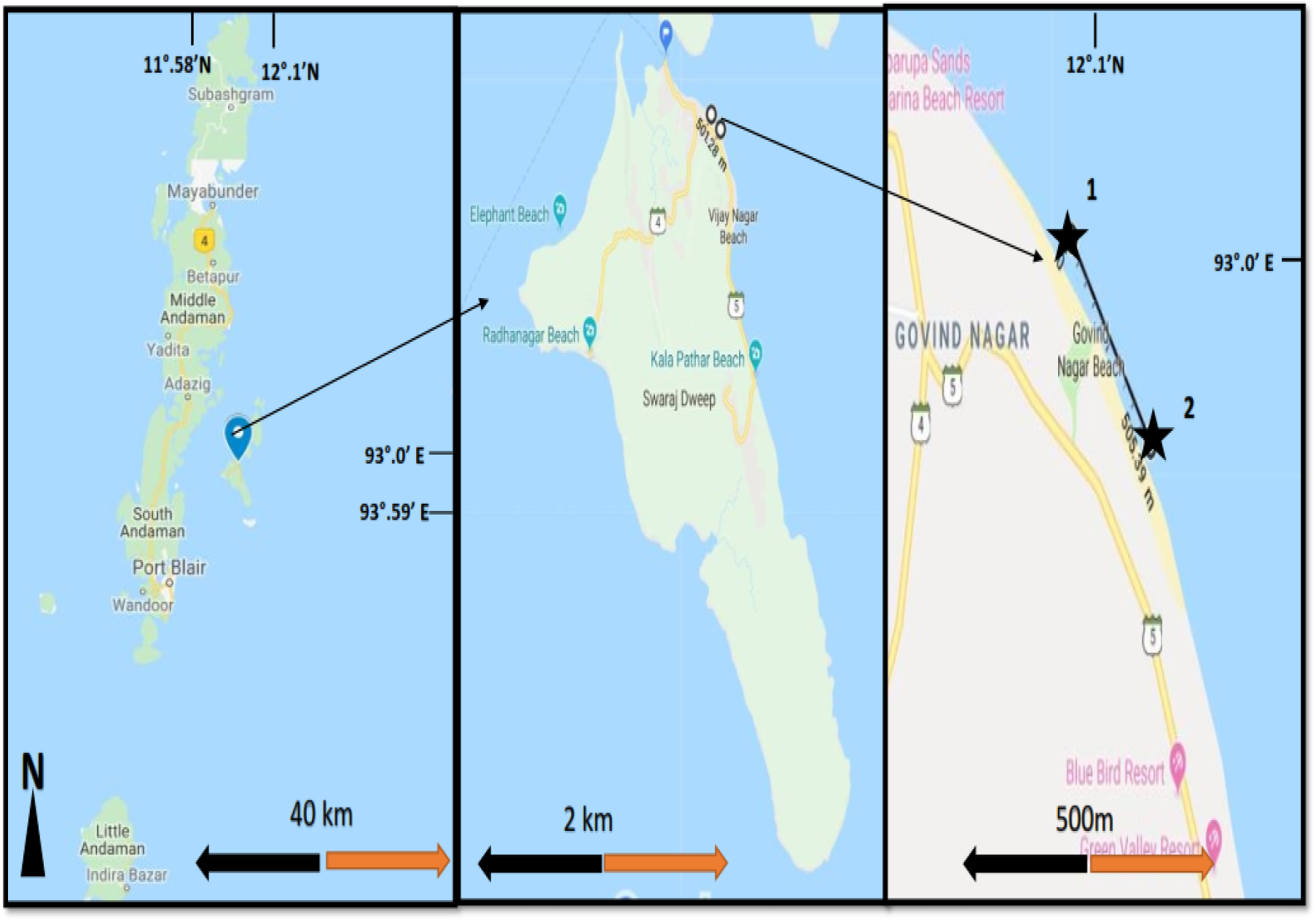
Study area showing station 1 and 2 of Govind Nagar beach of Havelock island in Andaman and Nicobar Islands

**Fig. 2.**
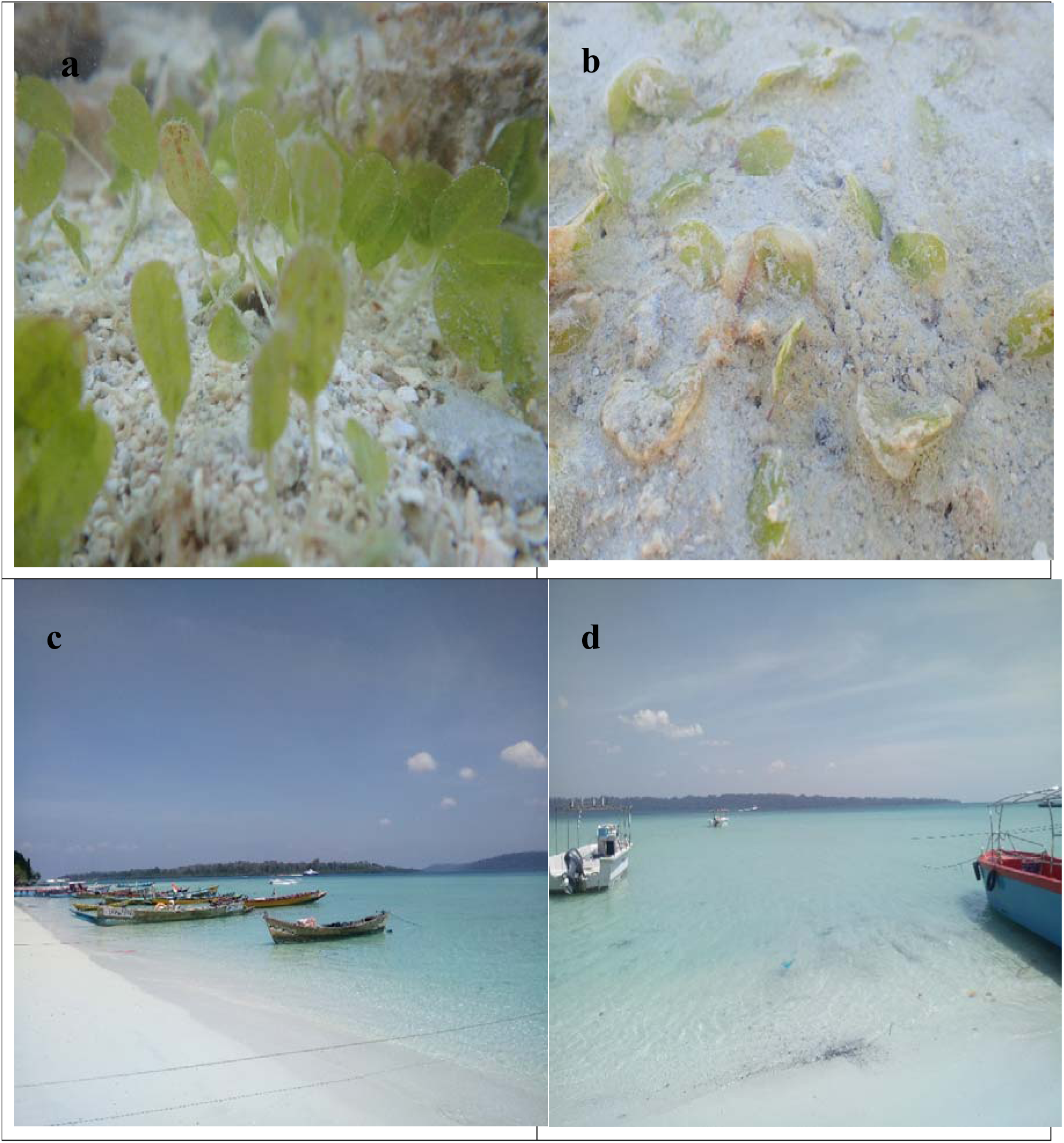
*H. ovalis* meadows recorded at a) sheltered area and b) exposed to anchoring and c) various kinds of fishing boats and d) recreational boats at station 1 used around Govind Nagar Beach of Havelock Island of Andaman and Nicobar Islands, India.

Sediment cores (n=5) were collected from each quadrat where seagrass was sampled using 5cm diameter and 10 cm long plastic core. Sediments were collected in plastic bags and were oven dried at 60°C for 72 hours before sieved for grain size fractions. We used a quadrat of 20cm^2^ and a hand shovel to dig out seagrass samples up to 10 cm depth in February-March of 2019. A clear sign of damage due to anchors was observed during the sampling. Quadrats (n=5) were collected from two separate stations of *H. ovalis* within a depth of 0.5m during low tide. From each quadrat seagrass leaves, rhizomes and roots were collected in plastic bags and brought to the lab for further analysis. Density was calculated by counting the number of shoots per each quadrat. Horizontal rhizome length, leaf length, width and height from shoot was measured using a Vernier Calliper. The canopy height of *H. ovalis*, i.e., the leaf length of the longest leaf from the sediment to the leaf tip was measured using a ruler (Mckenzie, 2007). After initial measurements the plant parts were separated and dried in a hot air oven at 60°C for 48 hours to get the dry weight biomass. The percentage cover of the seagrass was estimated from the area covered by seagrass from the total quadrat area.

One-way ANOVA was used to test the significant differences between *H. ovalis* density, biomass and morphometric features between the two stations. All data was pre-checked for normality and homogeneity of variance. Data were log transformed when normality and homogeneity of variance was not achieved for raw data. Data are presented as mean and standard error (S.E.).

## Results

Impact of boat anchoring on *H. ovalis* was found to be positive compared to sheltered areas. The sediment grain size analysis indicated 94 to 95% of the sediment as sand, whereas silt content was very low. The silt content at station 1 were 1.2-fold higher than station 2 (Table 1).

**Table 1.** Results of grain size analysis of sediments, biomass, morphometric, canopy height and percentage cover values of *Halophila ovalis* sampled from Havelock Islands of ANI, India. Mean ± Standard error (SE) values are presented. Small letters indicate significant differences between the two stations. Above ground (AB), Below ground (BG).

| Variables |  | Station 1 | Station 2 |
| --- | --- | --- | --- |
| Grain size (%) | Sand | 94.05 ± 5.55<sup>a</sup> | 95.25 ± 4.15<sup>a</sup> |
| | Silt | 5.95 ± 2.39<sup>a</sup> | 4.75 ± 2.09<sup>b</sup> |
| Biomass (g DW m<sup>-2</sup>) | AB | 0.71 ± 0.24<sup>a</sup> | 1.68 ± 0.23<sup>b</sup> |
| | BG | 2.72 ± 0.14<sup>a</sup> | 4.14 ± 0.79<sup>b</sup> |
| | AB:BG | 0.32 ± 0.04 | 0.57 ± 0.22 |
| No. of leaves shoot<sup>-1</sup> | | 5.87 ± 0.45<sup>a</sup> | 6.51 ± 0.87<sup>a</sup> |
| Canopy height (cm) | | 1.15 ± 0.04<sup>a</sup> | 2.20 ± 0.01<sup>b</sup> |
| Leaf length (cm) | | 1.02 ± 0.09<sup>a</sup> | 2.05 ± 0.09<sup>b</sup> |
| Leaf width (cm) | | 0.94 ± 0.01<sup>a</sup> | 1.23 ± 0.08<sup>b</sup> |
| Horizontal rhizome length (cm) | | 9.05 ± 0.08<sup>a</sup> | 15.26 ± 0.38<sup>b</sup> |
| Percentage cover (%) | | 20.3 ± 0.12 | 34.5 ± 0.89 |

Significant differences were observed for density (shoot and apex) of *H. ovalis* between station 1 and 2. The shoot density of station 1 (288.6± 14.8 no.m^-2^) was 1.2-fold lower than station 2(350.4± 60.2 no.m^-2^), whereas the apex density was similar between the two stations (Fig.3). The above ground and below ground biomass of *H. ovalis* were different and significant between the two stations. The above (0.71±0.24 g DW m^-2^) and below ground (2.72±0.14 g DW m^-2^) biomass of station 1 were 2-fold and 1.5-fold lower respectively, whereas the ratio of above ground to below ground (0.32±0.04) biomass was 1.7-fold lower than station 2(Table 1).

**Fig. 3.**
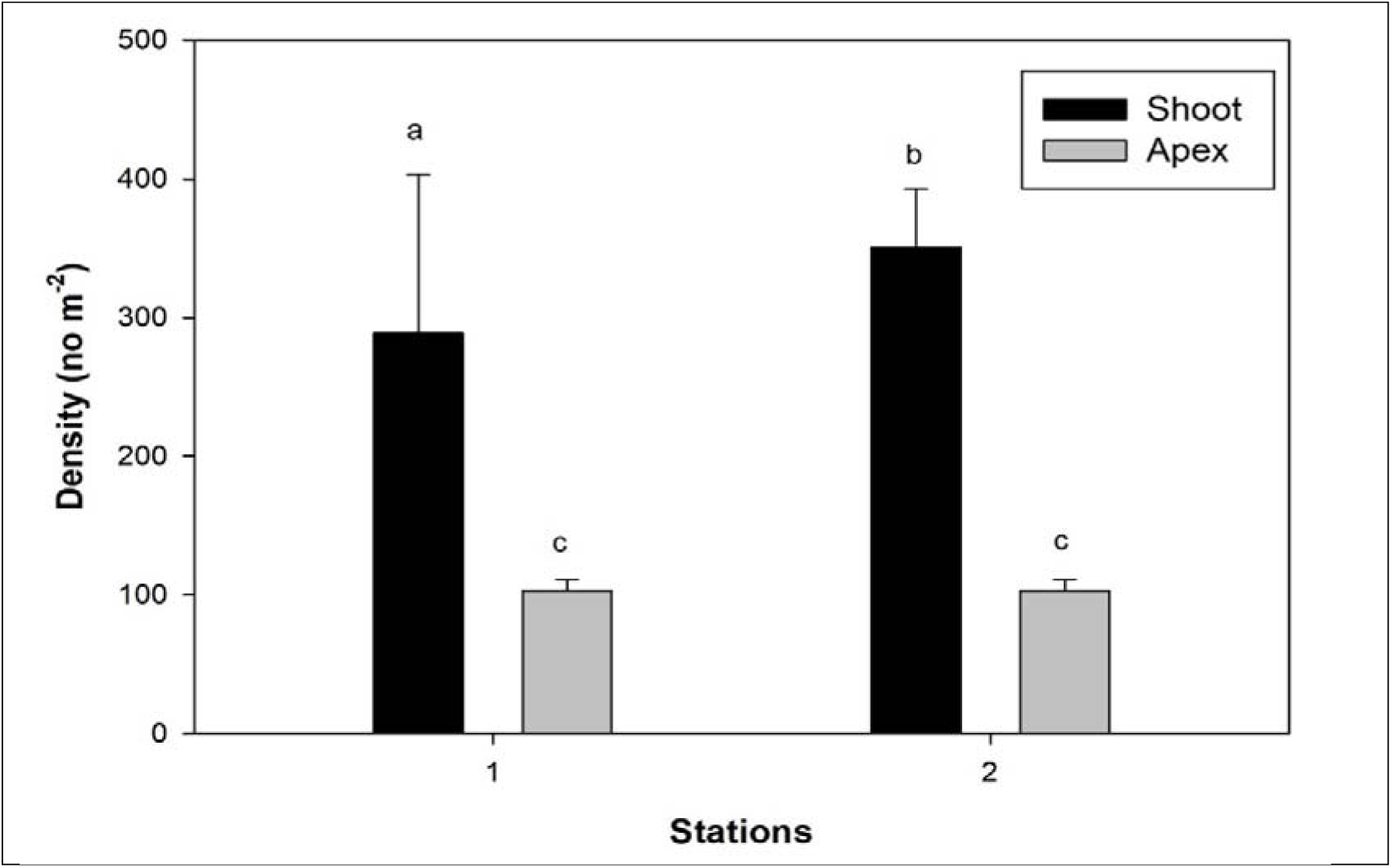
Density (Shoot and Apex) of *Halophila ovalis* of Havelock Island. Error bars represent standard errors. Significant differences between the two stations are represented by small letters

The morphometrics of *H. ovalis* were significantly different between the two stations, exception was number of leaves per shoot (Table 1). Among the two stations, the number of leaves per shoot was higher (6.51±0.87) at station 2. Both the leaf length (1.02±0.09 cm) and width (0.94 ± 0.01 cm) of *H. ovalis* were 2-fold and 1.3-fold lower respectively than station 2. The canopy height (1.15 ±0.04 cm) of seagrass at station 1 was 1.9-fold lower and the horizontal rhizome length (9.05 ±0.08 cm) was 1.6-fold lower than station 2. Reduction in morphometrics and density reduced the percentage cover (20.3± 0.12%) of *H. ovalis* at station 1, which was 1.6-fold lower than station 2 (Table 1).

## Discussion

Boat anchors causes disturbance to seagrass habitat that results in physical damage of individual plant structures and hampers the meadow growth and migration. In the current research we observed seagrass like *H. ovalis* patches has been greatly affected by continuous anchoring of boats that creates various scours interfering with the canopy of *H. ovalis* and their physical integrity resulting in loss of seagrass patches around Havelock Island, ANI, India. *H. ovalis* population structure were affected by boat anchors deployment compared to a sheltered station around the same site.

Sediment grain size of station 1 has a higher silt content compared to station 2 (Table 1), which can be the result of continues disturbance of the upper layer of the sediment by boat anchors, that induces the mobilization and dispersion of the impacted sediments by daily wave actions, along with crab holes (Unsworth et al., 2017). Higher silt content and softer sediments at station 1 can also contribute to the loss of shoot density as it increases the penetration of the anchors and can cause subsequent damage (Ceccherelli et al., 2007). Sediment erosion due to boat anchors not only lead to loss of *H. ovalis* patches but also contributes to loss of buried sediment organic carbon stocks and can change seagrass meadows as source of carbon rather than carbon sinks and contribute to increasing atmospheric carbon dioxide levels. Loss of seagrass meadows and subsequent loss of significant sediment organic carbon stocks been reported at Rottnest Island, Australia due to boat anchoring and mooring (Serrano et al., 2016).

Total density observed at station 1(391.7±11.7 shoots m^-2^) and station 2 (453.5±47.0 shoots m^-2^) were lower and similar (427.2±24.8) to the values reported for *H. ovalis* at Palk Bay, Tamilnadu (Gokulakrishnan and Ravikumar, 2014) and higher than previously reported density of *H. ovalis* from east coast of Malaysia (Sidik et al., 2010). Shoot density was significantly higher at station 2 than station 1 (Fig.2), which indicates the role of physical disturbances caused by boat anchors that damages the shoot structures of *H. ovalis*, which were evident during sampling. Loss of shoot density due to anchoring pressure was also reported for *Posidonia oceanica* in Mediterranean Sea (Francour et al 1999; Okudan et al., 2013).The apex density of *H. ovalis* at both stations were almost similar suggesting that higher sand (94-95%) and lower silt content in the sediment makes it difficult for the seagrass to migrate horizontally, as *H. ovalis* prefers higher silty habitat.

Density and biomass followed similar trends, i.e., lower density at station 1 were observed with lower above ground biomass and vice versa. Similar trends of lower density and lower above ground biomass was observed for *H. ovalis* of the east coast of Malaysia (Sidik et al., 2010). The above ground biomass of station 1(21%) and station 2 (29%) of the total biomass, were significantly lower for above ground biomass (42% of total biomass) reported for *H. ovalis* around the Andaman Sea (Erftemeijer and Stapel, 1999). However, these authors carried their research on deep water (1-16m depth) *H. ovalis*, which were more protected from the anthropogenic impacts directly. Saying that, lower above ground and higher below ground biomass in other seagrass species such as *Halophila beccarii* has also been observed around Andaman Sea (Aye et al., 2014). Higher below ground biomass of station 1 (79%) and station 2 (71%) coincide within the range of below ground biomass of 63-77% observed for *H. ovalis* meadows (Sidik et al., 2010) around Andaman Sea. Higher below ground biomass suggests that *H. ovalis* being smaller in size seagrass, needs extensive rhizome networks buried in the sediment for spatial meadow migration. Secondly fast-growing nature of *H. ovalis* can be the result of higher below ground biomass.

The number of leaves per shoot, leaf length and width of *H. ovalis* at station 1 were lower than station 2 (Table 1) because of the bending of leaf stem by the rope and anchors, resulting in breakage and burial of leaf structure in the upper layer of the sediment (Fig.2b) which inhibits the plant growth, productivity and biomass. However, there were not much difference in number of leaves per shoot between station1 and 2, because *H. ovalis* as a fast-growing tropical species can restore the leaves in quick time (Vermaat et al., 1995: Bharathi et al., 2014) damaged by boat anchors. Reduction in leaf length and width has been observed for other seagrass species like *P. oceanica* in Mediterranean Sea due to boat anchors (Montefalco et al., 2006).

The canopy height of *H. ovalis* at station 1 (1.15 ±0.04 cm) was lower than station 2 (2.20 ±0.01cm), which indicates about the physical injury the leaf structure sustains during drop down of boat anchors resulting in leaf scars and broken-down leaf-stems (Fig.2b) at station 1. While anchored the continuous swinging of the attached rope with the tidal movement, the size of the anchor and settlement of the boat during low tide on the seagrass leaves plays a significant role in determining the extent of damage. Once broken from stem the seagrass leaves are covered with sediments and other microbenthic algae that covers the seagrass leaf surface (Fig.2b), which alternatively reduces the seagrass photosynthetic capacity and its resilience to meadow development. Reduction of canopy height due to boat anchors were also observed for other seagrass species such as *Zostera marina* in UK (Unsworth et al., 2017), *P. oceanica* of Turkish coast, Mediterranean Sea (Okudan et al., 2013) and *Posidonia australis* in Australia (Demers et al., 2013). The observed canopy height of *H. ovalis* at both the stations were similar to canopy height of *H. beccarii* (0.7-1.5cm) observed at Kalegauk island (Aye et al., 2014), Myanmar and *H. ovalis*(1.98cm) of east coast of Malaysia (Sidik et al., 2010) in the Andaman Sea, where disturbances due to boat anchors have been reported.

The horizontal rhizome length being shorter at station 1 than station 2 clearly indicates the negative impact of physical damage of *H. ovalis* rhizome structure, which can result in meadow fragmentation and reduce the meadow migration of *H. ovalis*. Loss of rhizome structure and negative effects on meadow migration has been observed for climax species like *P. oceanica* in Mediterranean Sea (Francour et al., 1999). The overall negative impact of boat anchors on the morphometrics of *H. ovalis* resulted in low percentage cover, which has also been observed for other seagrass species like *Z. marina* in UK (Unsworth et al., 2017) and USA (Kelly et al., 2019), *P. oceanica* in Mediterranean Sea (Okudan et al., 2013; Ceccherelli et al., 2007; Montefalcone et al., 2006, 2008) and *P. australis* in Australia (Serrano et al., 2016)

The loss of seagrass patches from boat anchors at Havelock Island of ANI, India is small but significant at local scale as these disturbances lead to removal of *H. ovalis* biomass by shoot uprooting and breakage of leaves. These losses will impact the local biota that depend on *H. ovalis* meadows for food and habitat, such as *Dugong dugon*, which is already endangered mammal found in the waters of ANI and loss of his preferred feeding grounds can impact its conservation and recovery issues. Saying that, physical damages due to boat anchors fragments the seagrass meadows and reduces the collective adaptation and resilience of seagrass meadows to other anthropogenic stressors such as pollution and eutrophication (Unsworth et al.,2015; Maxwell et al., 2016). Loss of seagrass meadows will also reduce the extensive ecosystem services seagrasses provide, such as habitat for commercially important fish population and invertebrate biodiversity and carbon sequestration, (Macreadie et al., 2015; Cullen-Unsworth et al., 2018; Serrano et al., 2016).

We report for the first time about the effects of boat anchors on seagrass ecosystems of India. Our report will contribute to the status quo of *H. ovalis* patches of Havelock Island of ANI and to the development of National Status Report on Seagrass Ecosystems of India, an initiative from the Ministry of Environment, Forests and Climate Change, Govt. of India for better management and conservation practices for seagrass ecosystems under the influence of anthropogenic pressure.

## Conclusion

We find strong evidence that boat anchors cause loss of *H. ovalis* meadows. The loss of *H. ovalis* was restricted to the anchor deployment site with a clear indication of reduction in density, biomass, morphometrics, canopy height and percentage cover. Though this damage to seagrass meadows is local on small scale, it can lead to loss of feeding habitat for fish population and ecosystem services locally, which can have global impact. So, for proper management and conservation of seagrass ecosystems of India, the impact of habitat disturbances from boat anchors around the various seagrass ecosystems of India need to be reported.

## Acknowledgements

We are grateful to the PCCF, wildlife to provide the required official help during the sampling programme.

## Author Statement

AKM, SN has conceived the idea and have carried out the field work and data analysis. AKM has written the manuscript and SN and DA has provided valuable inputs to make the manuscript better. AKM, SN and DA give their full consent to publish the article.

